# Caloric restriction worsens decision-making impairments and gut dysbiosis after brain injury in male rats

**DOI:** 10.1101/2025.02.24.639886

**Authors:** Reagan L. Speas, Jenna E. McCloskey, Noah M. Bressler, Michelle A. Frankot, Carissa Gratzol, Kristen M. Pechacek, Kris M. Martens, Cole Vonder Haar

## Abstract

Traumatic brain injury (TBI) causes long-term deficits in decision-making and disrupts the gut microbiome. Dysbiosis of the gut microbiome is a potential contributor to the development of multiple psychiatric and neurological disorders and may be a contributor to chronic symptoms from TBI. Caloric restriction is often used to assess psychiatric-related behaviors in animals, but also affects the gut microbiome. In the current study, we evaluated the effects of caloric restriction versus free feeding on a frontal controlled cortical impact TBI. Rats were trained on the rodent gambling task, an analog of the Iowa gambling task, to assess risk-based decision-making. The microbiome was sampled through the acute to subacute period post-injury and lesion size and microglia counts evaluated at 10 weeks post-injury. Caloric restriction did not affect decision-making at baseline, but did affect motivational variables. TBI impaired decision-making and this effect was exacerbated by caloric restriction. Other motivation-related variables followed a similar pattern of impairment with TBI driving impairments that were worsened by caloric restriction. The gut microbiome was initially dysbiotic, but largely recovered within 14 days post-injury. Despite this, acute gut measurements were predictive of chronic decision-making impairment. These data indicate a role for the gut microbiome in the evolution of TBI deficits and suggest that interventions targeting the gut may have a limited window of opportunity to treat long-term deficits.

## Introduction

Traumatic brain injury (TBI) is a leading cause of death and disability with a global annual incidence rate of 27-50 million [1, 2]. TBI increases risk for psychiatric disorders as varied as attention deficit hyperactivity disorder and schizophrenia [3, 4]. Brain injury also contributes to a variety of psychiatric symptoms which may not fully meet the criteria of a disorder diagnosis, but nonetheless substantially affect quality of life [5]. One such symptom is risky decision-making, which affects everything from family and social relationships to financial well-being and risk for drug abuse. Recent evidence suggests that the gut microbiome is disrupted by TBI [6–8] and therefore, may be a contributor to developing psychiatric symptoms and disorders.

The gut microbiome is part of the gut-brain axis, a bidirectional network of communication linking the central nervous and the gastrointestinal systems to regulate digestion, influence peripheral organs, and modify neurological function [9]. The gut microbiome is composed of bacteria, archaea, viruses, and eukaryotes which exist in a balance to help support normal function. Various metrics of ecological diversity are used to quantify this balance of microbes, including measurements of alpha diversity (quantification of microbiome diversity within a subject) and beta diversity (quantification of dissimilarity between subjects). Disruption of the gut microbiome, measured by changes in diversity or direct measurements of specific bacteria, can directly affect neurological and psychiatric function [for an exhasutive review, see 10]. Such effects are well-established in depressive and anxiety-like behaviors as well as autism spectrum disorder. However, diseases involving impaired decision-making also implicate a role for the gut: bipolar disorder, attention deficit hyperactivity disorder, addictive disorders, obsessive compulsive disorder, and obesity [10]. Twenty-five years of studies on the gut-brain axis indicates that the microbiome should be a consideration when studying the etiology or development of psychiatric disease.

After TBI, there is high comorbidity with several of the same disorders linked to gut microbiome function [3, 4, 11]. A growing body of literature identifies gut dysbiosis after brain injury in clinical populations [12, 13] and animal models [6–8, 14]. Moreover, several recent studies demonstrated that manipulating the gut microbiome causally affected TBI outcomes. Specifically, fecal matter transplant worsened outcomes from TBI when they came from either epilepsy-prone rats or from a mouse model of Alzheimer’s disease [6, 15]. Further, antibiotic depletion/disruption of the microbiome mitigated some TBI inflammatory pathology [16]. Despite these findings, relatively little work is being done to evaluate how psychiatric-related outcomes from TBI could be linked to the gut microbiome. A recent clinical sample of veterans with a history of blast exposure showed some minor associations of microbial measures with executive functions and depressive measures [17]. However, the heterogeneous and highly-comorbid nature of the sample made strong inference with regard to TBI difficult. In a prior paper using a rat model of TBI, we established that chronic decision-making deficits from TBI were predicted by acute gut diversity measurements [8]. These limited samples indicate that further study of the relationship between psychiatric-related functions, brain injury, and the gut microbiome is needed. Animal models provide a strong method to isolate both the effects of injury on the gut and the effects of the gut microbiome on the injury process. However, one potential limitation in studying psychiatric-related outcomes in animal models is that they largely rely on motivated behavior, and these behaviors are primarily motivated by food restriction.

Mild food restriction is often used as the “establishing operation” to increase motivation to respond for food-based reinforcers [18]. However, caloric restriction also has effects on the gut microbiome. Broadly, these effects appear beneficial: helpful taxa such as *Lactobacillus* were increased [19], however, potentially detrimental taxa such as Proteobacteria also increased with larger caloric restriction [19, 20]. Caloric restriction can decrease gut diversity, prime the gut for opportunistic infection [21], and increase expression of tight-junction proteins to restrict bacterial passage across the intestinal wall [19]. Historically, our lab group and others use mild food restriction to motivate behaviors with little regard to how this might interact with the gut microbiome. This form of caloric restriction has not been evaluated to see how it influences post-TBI psychiatric-like symptoms, potentially via the gut microbiome. A method using mild food restriction to capture deficits in decision-making after TBI is the Rodent Gambling Task (RGT), a measure of decision-making under risk [22]. In our research, brain injury reduces optimal decision-making and shifts preference towards both suboptimal and riskier options [23, 24]. Gut microbiome dysfunction after TBI is predictive of long-term impairment in this behavior [8], but it is not clear if that was influenced by caloric restriction.

The goal of the present study was to evaluate how caloric restriction to approximately 85% free feeding weight would affect rats’ choice on the RGT, affect the outcome of a frontal TBI, and alter the gut microbiome. We hypothesized that caloric restriction would affect decision-making on the RGT as well as motivational measures, but that the effect of TBI would be similar on behavior and the gut microbiome. We also used exploratory regression models to understand how the gut microbiome may directly relate to progression of TBI deficits. Data from this could then be used to inform a better understanding of potential mechanisms by which the gut microbiome may shape TBI outcomes and lead to novel therapeutic avenues for decision-making impairments after TBI.

## Materials and Methods

### Experimental Design

In a two-group design, we placed male rats into different feeding conditions (Restricted vs. Free-fed). We then evaluated their acquisition of probability-based decision-making on a rat analog of the Iowa Gambling Task as previously described [22]. This allowed an assessment of feeding condition on acquisition of decision-making behavior. After choice behavior stabilized, both groups received a frontal TBI, allowing group-level comparisons for the effect of feeding condition and within-subject pre- versus post-behavior comparison for the effect of injury. After 7 weeks of testing, we reversed the feeding conditions to evaluate long-term versus acute effects of caloric restriction on behavior. We collected fecal samples to evaluate the gut microbiome at time points of pre-injury and 1-, 3-, 7-, and 14-days post-injury. The timeline is summarized in Figure 1A.

**Figure 1.**
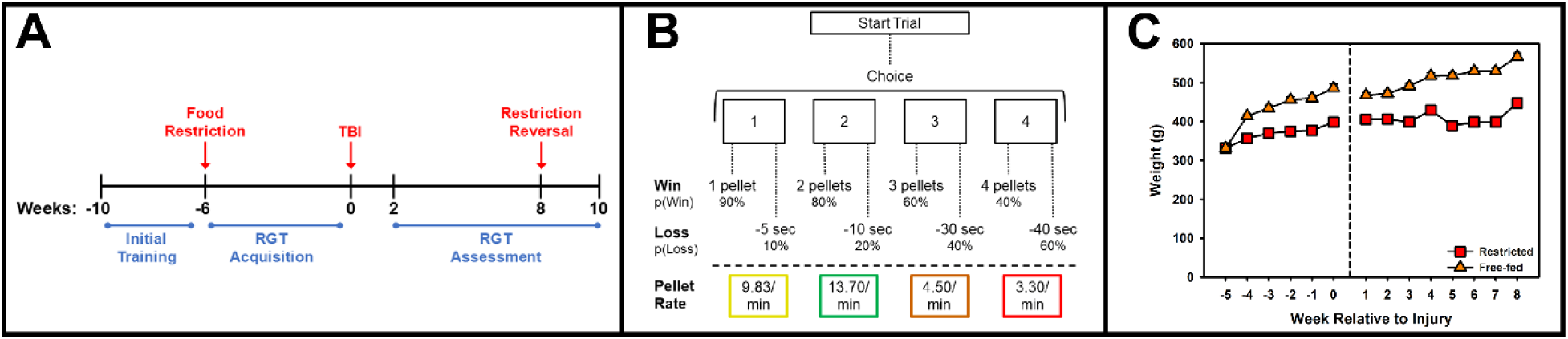
Experimental design and task layout. Rats were pseudorandomly assigned to Restricted or Free-fed condition prior to training on the Rodent Gambling Task (RGT), then given a TBI after acquisition of the behavior (panel A). They were then re-assessed from weeks 2-8 post-injury before reversing the feeding conditions and then euthanized at 10 weeks post-injury. Microbiome samples were collected prior to injury and at 1-, 3-, 7-, and 14-days post-injury. The RGT task (panel B) provided four choice options, each with varying probabilities of reinforcement (“win”) and punishment (“loss”), leading to an optimal (2-pellet option), suboptimal (1-pellet option), and two risky (3- and 4-pellet options). (C) Rats assigned to the food restriction condition weighed 80-85% less than rats assigned to the free-fed condition.

### Animals

Subjects were 24 male Long-Evans rats (Charles River, Wilmington, MA). Subjects were 2 months old at the beginning of training, 4 months old at point of injury, and 6 months old at time of euthanasia. Rats were matched for performance as measured by sessions to acquire the nose-poke response and pseudorandomized into free fed (*n =* 12, “Free-fed”) or calorie restricted (*n =* 12, “Restricted”) groups. The Restricted rats were maintained at 14 g chow per day plus any pellets earned, and the Free-fed rats consumed approximately 20-22 g chow per day plus any pellets earned. Weights were taken weekly. Subjects were pair-housed prior to surgery and single-housed after surgery on a reverse light cycle. One subject was excluded at euthanasia due to a lack of brain injury.

### Behavioral Apparatus

Behavioral testing took place in a standard five-hole operant conditioning chamber with a stimulus light in each recessed hole (Med Associates, St. Albans, VT). Nose pokes were measured with breaks to an infrared beam in each hole. A food hopper with a light was located on the opposite wall with a pellet dispenser above it. Nutritionally-complete purified sucrose pellets were used as reinforcers (45 mg F0021, BioServ, Fleming, NJ), and behavioral tests were programmed using software written in Med-PC IV.

### Pre-Injury Rodent Gambling Task (RGT) Acquisition

Rats were trained as previously reported [22, 23]. Training began with a 30-min habituation session, where sucrose pellets were placed in each of the five holes and in the food hopper. Once habituated (consuming all pellets in a session), rats were trained and evaluated on the initial stages of the five-choice serial reaction time task [25] to condition a response to a lit hole. Rats initiated trials with a nose-poke into the food hopper, which prompted a 5 s delay before illuminating one of the five holes. Correct responses were reinforced with one sucrose pellet. The time to respond to the cued hole was 30 s, which was reduced to 20 and 10 s over sessions as the rat accurately responded within the time limit. Premature, omitted, or incorrect responses were punished with a 5 s timeout with the house light illuminated. Sessions lasted 30 min or a maximum of 100 total trials. Training continued until each rat completed at least 50 trials with 80% accuracy with the 10 s stimulus. Rats were matched for performance on acquisition and then randomized into Free-fed or Restricted groups. They then began RGT training.

During the RGT, rats had 30 min in a session to maximize reinforcement (Fig 1B). Four choice options were given with a varying reinforcement (sucrose pellet) and punishment (timeout) probabilities and magnitudes. Choice P1 had a 90% chance of earning one sucrose pellet and a 10% chance of a 5 s timeout. Choice P2 had an 80% chance of earning two sucrose pellets and a 20% chance of a 10 s timeout. Choice P3 had a 60% chance of earning three sucrose pellets and a 40% chance of a 30 s timeout. Choice P4 had a 40% chance of earning four pellets and a 60% chance of a 40 s timeout. Initially, rats were trained on a forced-choice version of the RGT, where they were given seven, 30-min sessions to expose them to the consequences associated with each choice. Once completed, rats were allowed to make free choices for the remainder of the study. During each trial, a nose-poke to the food hopper would trigger a 5-s delay (intertrial interval). A response during this period was considered “premature” and would initiate a 5-s timeout with no opportunity to earn reinforcement. All choice holes illuminated after the intertrial interval and the rat was given a 10 s limited hold to respond. After making a choice, either the hopper would light up and the associated magnitude of sucrose pellets would be delivered, or the associated punishment timeout would be initiated, with the choice hole flashing at 1 Hz for the duration. Omitted choices were punished with a 5-s timeout. Hole location for each choice was maintained throughout the study and counterbalanced with two analogous forms of the program. Rats reached a stable baseline (determined by visual inspection at subject level) after 29 free-choice sessions and advanced to surgery.

Multiple variables of interest can be extracted from the RGT. The primary is the percent choice amongst the different options: P2, the optimal choice; P1, the suboptimal choice, P3 and P4, the two risky choices (Fig 1B). Subsequent variables can be derived from this choice, including likelihood to stay with the same choice after “wins” or “losses”, and a “score” variable that encompasses all four options ([P1+P2] – [P3+P4]). Premature or impulsive responses are also of interest as are various motivational variables such as latencies to choose or collect reinforcers as well as omitted trials. The total pellets earned per session can also be calculated which encompasses all variables together.

### TBI Surgery: Controlled Cortical Impact

All subjects received a bilateral, frontal TBI via controlled cortical impact (CCI) procedure as previously described [23, 26]. Briefly, rats were anesthetized with isoflurane (5% initially, 2-4% maintenance in 0.5 L/min oxygen). Rats were placed in a stereotax. At the incision site on the skull, local anesthesia (0.1 ml bupivacaine, 0.25%, s.c.) was given and on the body, general analgesia was given (carprofen, 5 mg/kg, s.c.). The incision site was made aseptic, a midline incision was performed, and periosteum removed. A circular 6 mm craniectomy was performed with a surgical drill, centered on the midline, 3.0 mm, in front of bregma. A moderate-severe, bilateral, frontal CCI was generated using an electromagnetic impactor (Neuroscience Tools , O’Fallon, MO). The injury was made with a 5 mm diameter flat-faced tip, to a depth of 2.5 mm, and at a velocity of 3 m/s with a 0.5 s dwell time. Once bleeding ceased, the incision was sutured and antibiotic ointment administered at the incision. The TBI surgeries took approximately 30 min each. Post-surgical care was given for seven days, which included daily monitoring, free feeding, and analgesia as needed.

### Post-Injury Rodent Gambling Task (RGT) Assessment

After seven days of recovery, rats were re-tested on the RGT. Testing continued five days per week until 10 weeks post-injury. During Weeks 9 and 10, the feeding conditions were reversed as described above.

### 16S rRNA Sequencing of Gut Microbiome

Fecal samples were collected prior to injury, and at 1, 3, 7, and 14 days post-injury into a clean microcentrifuge tube on ice. Samples were collected directly from the anus when available in the morning or, when unavailable, the rat was placed in a fresh cage and sample taken later that day from the rat or cage. Samples were stored at -20 °C until DNA extraction. Fecal samples were processed using a kit (PureLink Microbiome DNA, #A29790) and DNA concentration validated using a spectrophotometer (NanoDrop 2000, ThermoFisher Scientific, Waltham, MA). Samples were sent for 16S rRNA sequencing (Novogene, Sacremento CA). Samples were amplified using the V3-V4 region of the bacterial 16S gene.

Demultiplexed sequenced data containing the pair-ended samples were imported into R statistical software and processed using the *dada2* package [27]. Samples were filtered by removing primers and low-quality reads. The dada2 algorithm was used to cluster sequences into unique amplicon sequence variants (ASVs), merge paired reads, and remove chimeras.

ASVs were compared against a known taxonomy list from the Silva Project [28]. This resulted in an ASV table that contained the frequency of the unique variants within each sample at the level of the kingdom, phylum, class, order, family, and genus. Phyla are referred to by common names: Firmicutes (now Bacillota), Proteobacteria (now Pseudomonadota), Bacteroidota (formerly Bacteroidetes).

### Histology

At the conclusion of behavior, rats were transcardially perfused with 0.1M PBS, followed by 3.7% formaldehyde. The brains were extracted and post-fixed with 3.7% formaldehyde for 24 hours before prior to storage in a 30% sucrose + 0.02% sodium azide solution. Brains were frozen and section at 30 µm using a sliding microtome. Slices were then selected for immunohistochemistry or mounted on glass slides for Nissl staining. Nissl staining was performed with thionin to measure remaining brain volume. Briefly, slides were rehydrated using a series of washes of decreasing ethanol concentrations, placed in thionin solution, then dehydrated with a reverse sequence. Slides were then cover slipped and dried. The slides were digitally scanned (600 DPI) and brain slices surrounding the lesion (+1.0, +2.0, +3.0, +4.0, and +5.0 mm from bregma) were measured using ImageJ (NIH, Bethesda, MD). Brain volumes were estimated by multiplying the average size of the slices by their thickness (30 um).

IBA-1 staining was used to visualize microglia in the peri-lesion and hippocampus regions using a free-floating method. Briefly, slices were blocked with peroxidase, rinsed, then incubated in 2% Normal Goat Serum overnight (4° C). After each of the following steps, slices were rinsed in PBS+0.1% Tween, then PBS. Slices were incubated in primary rabbit anti-IBA-1 (1:2000, Wako 019-19741) for 48 h at 4° C, followed by goat anti-rabbit IgG (1:2000, Vector Laboratories BA-1000) for 90 min at room temperature. They were then placed in Vectastain ABC solution (Vector Laboratories PK-4000) for 90 min at room temperature, followed by a reaction with a 3-3’-diaminobenzidine peroxide solution (0.05% DAB & 0.015% H_2_O_2_ in PBS). Slices were then mounted, dehydrated, and cover slipped for imaging. The brains were imaged at 40x magnification using a Fisherbrand Micromaster microscope and Swiftcam 16MP camera. Microglia were counted automatically using ImageJ. The phenotype of the microglia were analyzed using ImageJ’s automated circularity index (0 = no circularity, 1 = perfect circle) and evaluated at four different circularity settings (0, 0.1, 0.2, 0.3) to measure potential phenotypic shifts.

## Data Processing and Analysis

All data were analyzed using R statistical software (http://www.r-project.org/; version 4.4.1) with the *brms*, *lme4*, *lmerTest, emmeans*, *vegan*, and *stats* libraries. The critical *p*-value was set to 0.05. Most behavioral data on the RGT were analyzed in linear mixed effects regression (LMER). Transformations (square root, log, arcsine-square root) were applied as needed to normalize data. The following variables were analyzed: percent premature responses, percent omitted responses, total trials, pellets earned, choice latency, reinforcer collection latency, choice score, and likelihood to stay with a choice given a win or loss.

Proportional choice data were transformed with an arcsine-square root transformation and analyzed with a Bayesian LMER where choice proportions were allowed to vary as a function of subject. We have reported this method to be less sensitive to false positives for compositional/interdependent data [29]. Estimated marginal means and slopes were used to compare significant effects and interactions.

Microbiome data were merged with study variables and alpha (Shannon index, Inverse Simpson index, Observed OTUs) and beta (Bray-Curtis dissimilarity) diversity were calculated using the *phyloseq* library [30]. The effect of feeding condition on pre-injury alpha diversity was analyzed using Students *t*-tests. After injury, the Condition x Time effect on alpha diversity was analyzed using LMERs, where baseline alpha was included as a covariate and allowed to vary by subject. Estimated marginal means were used to compare significant effects and interactions. The multivariate outcomes beta diversity and taxa abundance (minimum 1% abundance) were analyzed using PERMANOVA in the *vegan* library [31], assessing the Condition effect pre-injury and the Condition x Time effect post-injury. A Wilcoxon paired-rank post hoc test was used when main effects were statistically significant.

A series of regression models were conducted to evaluate potential contributions of the microbiome to behavioral outcomes. The choice of the optimal P2 option served as the dependent variable. Both mixed effects (i.e., baseline behavior allowed to vary by subject) and fixed-effects only models were used to counteract issues with overfitting to the individual subject. Models were compared on the Akaike Information Criterion (AIC) and Bayesian Information Criterion (BIC) with the lowest values selected as the best-fitting model. First, we assessed stable pre-injury choice of the optimal P2 option as a function of pre-injury microbiome measurements. This allowed interpretation of effects of the microbiome on choice behavior in the absence of injury. Alpha diversity (summary level) as a predictor was compared against additive models of phyla abundance (Firmicutes, Bacteroidota, Proteobacteria) and Firmicutes/Bacteroidota ratio. Second, we assessed choice of optimal P2 at post-injury behavioral stability (final two weeks prior to condition reversal) as a function of individual microbiome measurements taken at the different time points (pre, 1D, 3D, 7D, 14D post-injury). This allowed us to evaluate the degree to which these measurements correlated to behavior, but did not account for the injury or feeding condition. Alpha diversity, phyla abundance, and Firmicutes/Bacteroidota ratio were used as predictors in separate models. Finally, we assessed the same chronic post-injury outcome as a function of microbiome measurements with additive predictors of Condition and Baseline Choice. This allowed us to determine whether microbiome measurements offered additional predictive power beyond just knowing the feeding condition and injury effect (change from baseline).

Postmortem histology was analyzed with a Student’s t-test (brain volume) or generalized LMER with a Poisson distribution (IBA-1).

## Results

Full tables of all results can be found in the supplement, including random effect descriptions. Here we present only key values and comparisons. Analysis syntax will be made available on the author’s GitHub.

### Rat Weights

Restricted rats had reduced weight at approximately 80-85% of the Free-fed condition (Fig 1C).

### Pre-Injury RGT Performance

To evaluate the primary outcome of the RGT, percent choice of each option, we performed a Bayesian LMER of acquisition (fixed effects: Percent Choice ∼ Condition*Week; random effects: Choice Option by Subject) and after stable baseline emerged (final week of testing). There was a significant Condition by Week interaction, such that the slope for all four options differed across feeding Condition (β: 0.06, CI: 0.02 to 0.10). Specifically, Free-fed rats increased rate of choice of the optimal P2 option and decreased rate choice of the suboptimal P1 option compared to risky P3 and P4 options. However, after choice stabilized, there was no significant difference between Condition (β: 0.16, CI: -0.35 to 0.68). Figure 2 (left of line break) displays the acquisition choice data.

**Figure 2.**
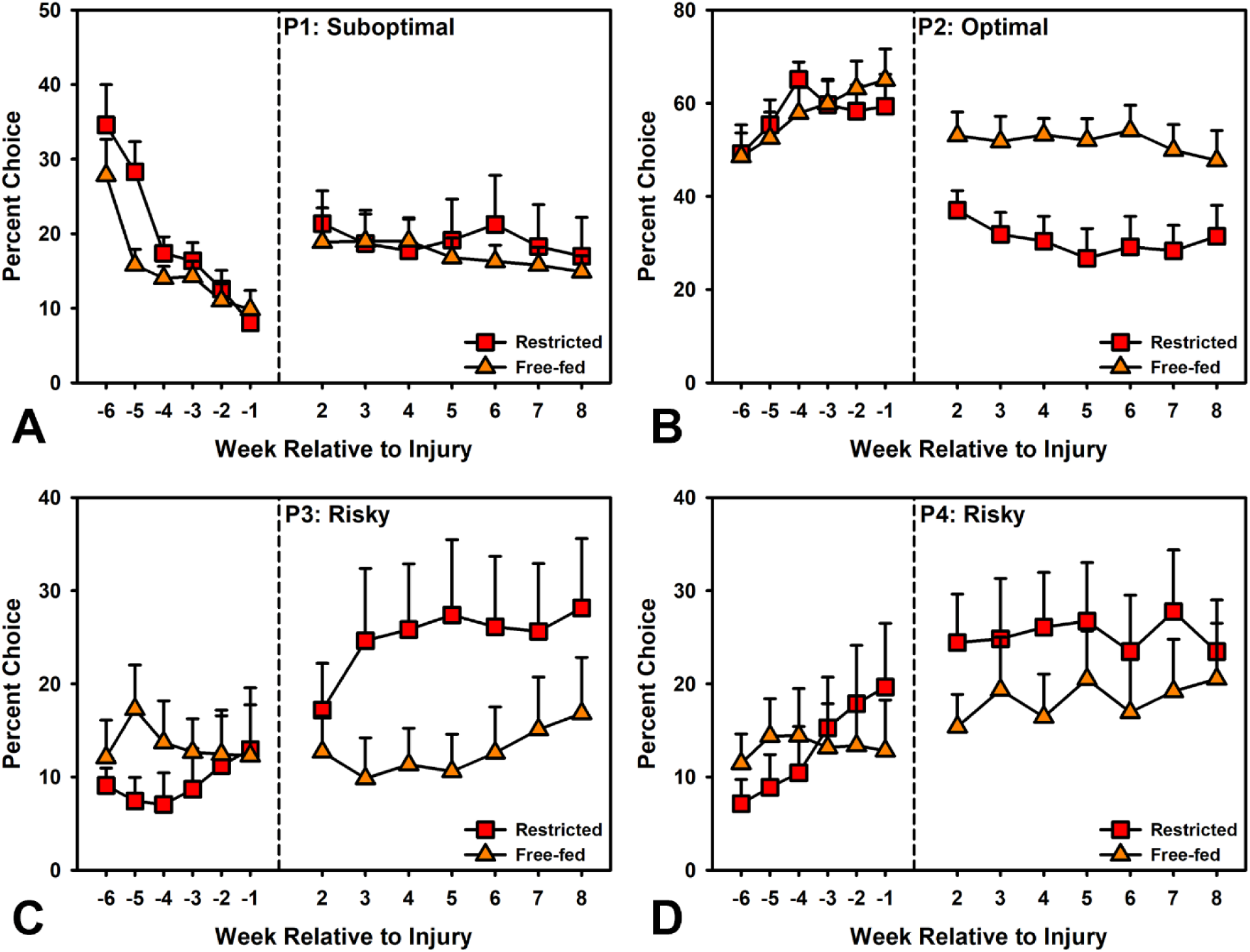
Acquisition and Post-Injury RGT Decision-making Outcomes. There was a significant Condition x Week interaction such that the Free-fed group acquired preferences across all options slightly differently (panels A-D, left of break; estimate CI: 0.02 to 0.10). However, there were no differences at baseline (-1 weeks relative to injury; estimate CI: -0.35 to 0.68). After injury (right of break), the Restricted group performed worse on the P2 optimal option (panel B; estimate CI: -1.23 to -0.27) but was not significantly different on the others. A separate within-subject analysis confirmed an effect of injury as measured by change from baseline for both groups across all options, with the exception of the Free-fed group on the P3 Risky option (panel C).

We also evaluated the effect of feeding condition on other RGT variables using LMER (fixed effects: Outcome ∼ Condition*Week; random effects: Subject intercept). Several variables were significantly different across acquisition of the task (see Supplement). However, at baseline, the only variables that remained significantly affected by Condition were those dealing with task motivation (Omissions, Trials completed, Pellets earned, Latencies; *p*’s < 0.009) and impulsivity (Prematures; *p* = 0.001). Specifically, the Free-fed rats completed fewer trials, made more omissions and therefore earned fewer reinforcers. They were also slower to respond in the choice phase and collect the reinforcer. They also made fewer impulsive responses. Figure 3 (left of line break) displays the acquisition data.

**Figure 3.**
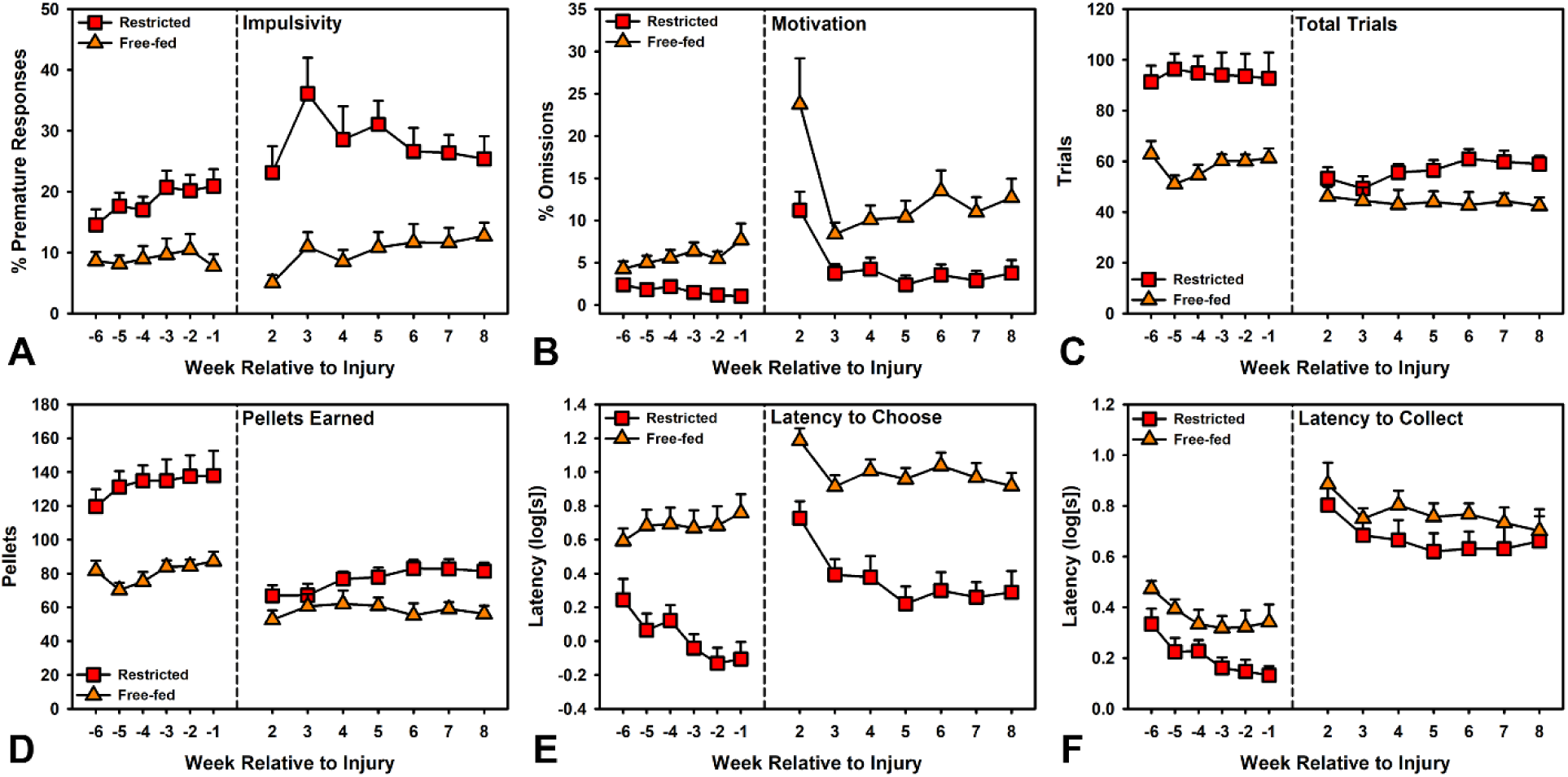
Acquisition and Post-Injury RGT Other Variables. During task acquisition (left of break) and at baseline (-1 weeks relative to injury), Free-feeding significantly altered variables measuring Impulsivity (panel A) and aspects of motivation (panels B-F; *p*’s < 0.05). After injury (right of break), these group differences persisted except for Collection latency (panel F; *p* = 0.149). A separate within-subject analysis confirmed there was still an injury effect across all variables for both groups on the motivational variables (panels B-F; *p*’s < 0.003). However, there was no injury effect on impulsivity for the Free-fed group (panel A; *p* = 0.719).

### Post-Injury RGT Performance

After injury, we evaluated the effect of feeding Condition on percent choice of each outcome on the RGT using a Bayesian LMER (fixed effects: Percent Choice ∼ Condition*Week + Baseline; random effects: Choice Option by Subject). There was a significant difference between the Conditions (β: 0.65, CI: 0.13 to 1.14). Specifically, the Restricted rats had a larger drop in preference for the optimal P2 option, but the relative change in P3 and P4 preference was not significant. Because there was no sham control, an additional analysis was performed to evaluate change from baseline to confirm the injury affected choice outcomes for both groups (fixed effects: Percent Choice ∼ Phase; random effects: Choice Option by Subject). Injury significantly changed choice preference from baseline for both Restricted (β: -0.94, CI: -1.10 to - 0.78) and Free-fed groups (β: -0.45, CI: -0.59 to -0.31). Restricted rats significantly changed preference from optimal P2 to suboptimal P1, risky P3, and risky P4 while the Free-fed rats changed preference from optimal P2 to suboptimal P1 and risky P4. Figure 2 (right of line break) displays the post-injury data.

The other RGT variables were similarly evaluated using LMER for Condition (fixed effects: Outcome ∼ Condition*Week + Baseline; random effects: Subject intercept) and to confirm injury effect (fixed effects: Outcome ∼ Phase; random effects: Subject intercept). The significant differences between the groups on most motivational variables remained the same (Omissions, Trials completed, Pellets earned, Choice latency; *p*’s < 0.030), however there was no longer a difference in reinforcer Collection latency (*p*’s > 0.149). Premature/impulsive responses also remained significantly lower in the Free-fed condition (*p* < 0.001). Despite the differences in Condition, an injury effect was observed across all variables for both groups (*p*’s < 0.050) except premature/impulsive responses for the Free-fed rats (*p* = 0.719). Figure 3 (right of line break) displays the post-injury data.

### Effects of Reversal of Feeding Conditions on RGT Performance

To dissociate which effects resulted from chronic mild caloric restriction versus acute hunger, we reversed the feeding conditions in Weeks 9 and 10 post-injury to obtain 8 sessions of data. Data were evaluated in Bayesian LMER for Choice data (fixed effects: Percent Choice ∼ Condition*Session + Baseline; random effects: Choice Option by Subject) and LMER for other variables (fixed effects: Outcome ∼ Condition*Session + Baseline; random effects: Subject intercept). Switching the feeding condition did not affect RGT choice (β: 0.05, CI: -0.14 to 0.24; Fig 4). However, it did affect all other motivational variables (Omissions, Trials completed, Pellets earned, Collection latency; *p*’s < 0.001; Fig 5) except Choice latency (*p* = 0.108; Fig 5E). Premature/impulsive responses were also reversed with the condition change (*p* = 0.002; Fig 5A).

**Figure 4.**
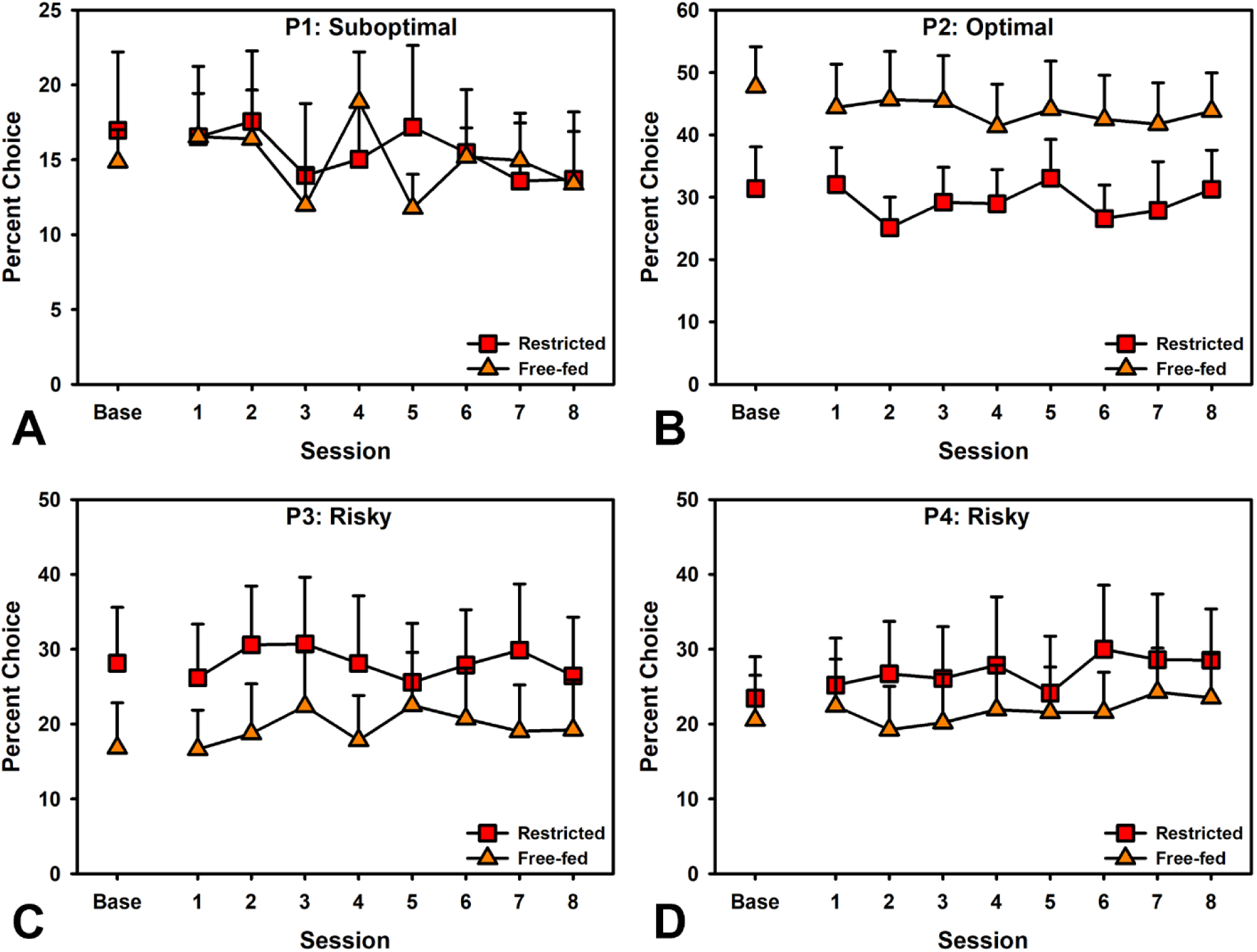
Effects of Feeding Condition Reversal on Decision-making. Reversing the feeding conditions had no effect on decision-making in the RGT (estimate CI: -0.14 to 0.24). Note: key label refers to *originally-assigned condition* at time of injury.

**Figure 5.**
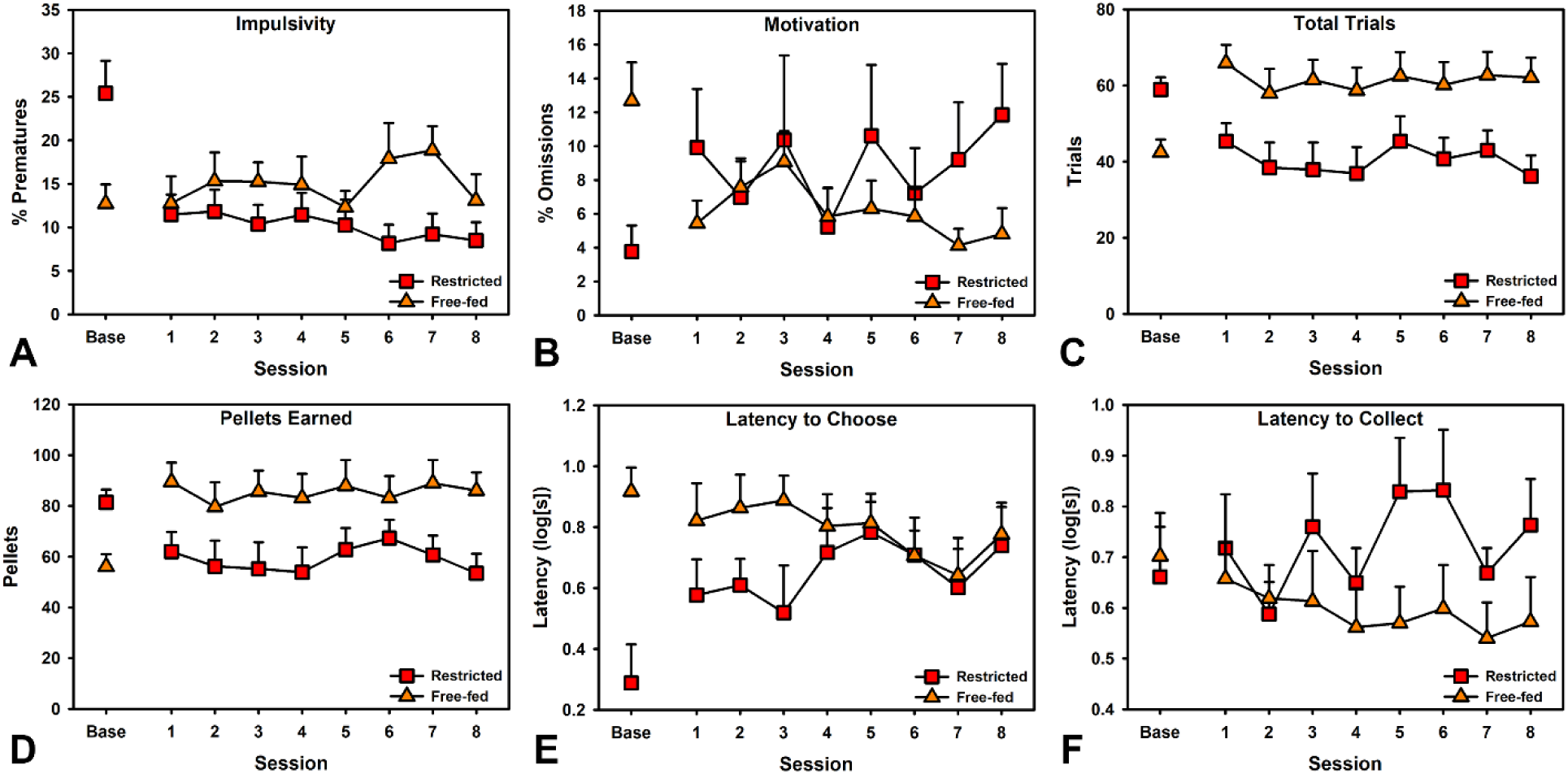
Effects of Feeding Condition Reversal on other RGT Variables. Reversing the feeding conditions significantly affected Impulsivity (panel A; *p* = 0.002) as well as the motivational variables Omissions, Trials completed, Pellets earned, and Collection latency (panels B-D, F; *p*’s < 0.001). However, there was no effect on Choice latency (panel E; *p* = 0.108). Note: key label refers to *originally-assigned condition* at time of injury.

### Pre-Injury Microbiome Analysis

Pre-injury differences in alpha diversity were analyzed by t-test, while beta diversity and phyla composition were analyzed by PERMANOVA. For alpha diversity, there were no significant effects of Condition on the Shannon index, Inverse Simpson index, or observed OTUs (*p*’s > 0.293). There were significant differences by Condition in beta diversity as measured by the Bray-Curtis metric (*p* = 0.005). However, when individual phyla were analyzed, there was no difference in proportions (*p* = 0.419). These data are shown in Fig 6.

**Figure 6.**
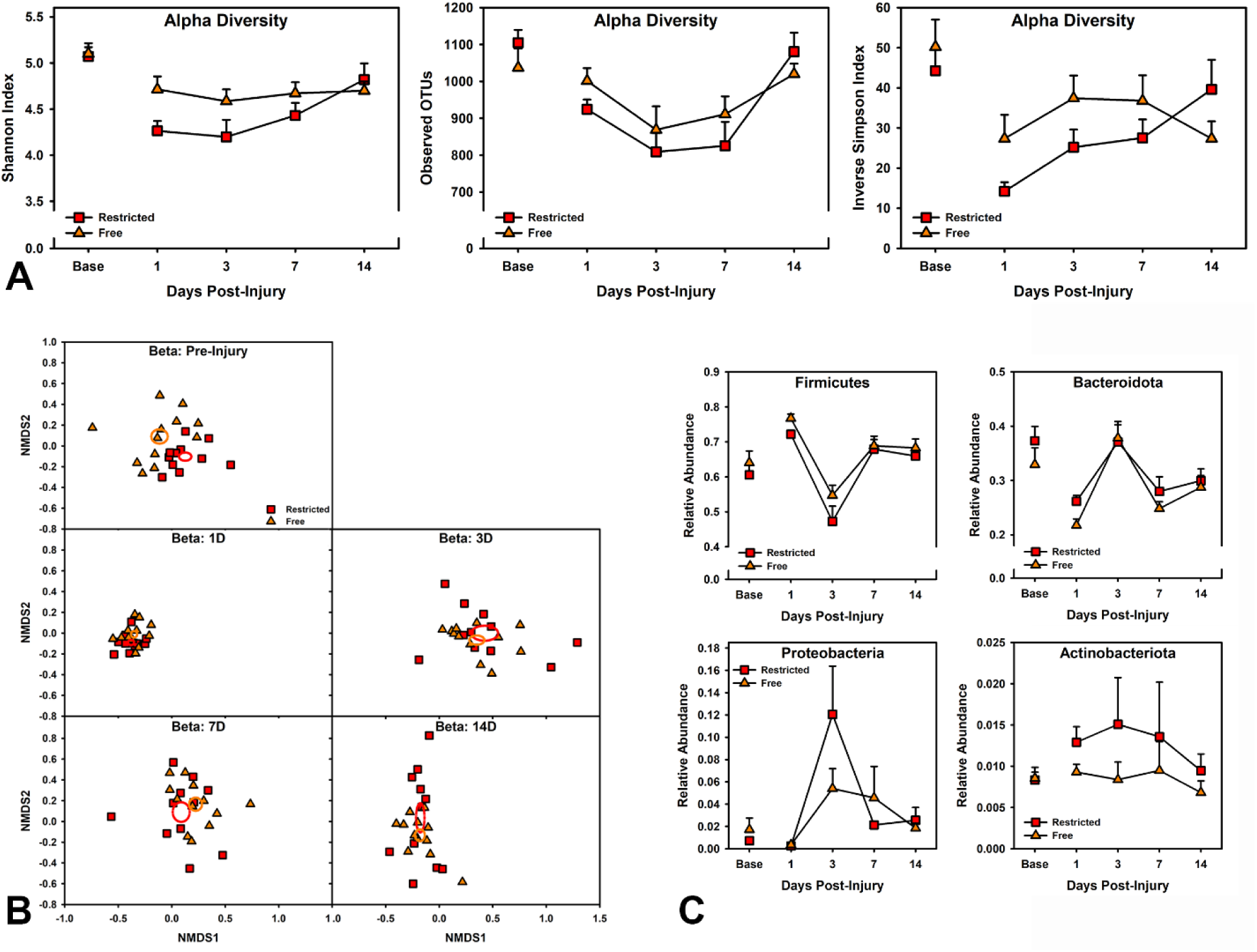
Gut Microbiome Changes due to TBI and Feeding Condition. At the pre-injury baseline, there were some microbiome differences in beta diversity (panel B; *p* = 0.005), but not alpha diversity or taxa abundance (panels A, C). For alpha diversity after injury (panel A), the Shannon index indicated a significant decrease in the Restricted condition (*p* = 0.030), while the observed OTUs indicated change in both groups by a main effect of Time (*p* < 0.001), and Inverse Simpson index indicated a Condition X Time interaction (*p* = 0.045). Considered together, these appear to be a main effect of injury with a potential caloric restriction interaction further reducing diversity. For beta diversity (panel B), the conditions were significantly different (*p* = 0.007) and changed with time (*p* = 0.001), but the interaction was not significant (*p* = 0.051). In the observed taxa (panel C), there were no significant group differences (*p* = 0.075), but a main effect of Time (*p* = 0.001), indicating an overall injury effect as phyla shifted across the measured days.

### Post-Injury Microbiome Analysis

Alpha diversity metrics were analyzed with LMER (fixed effects: Condition*Time[categorical] + Baseline, random effects: Subject intercept). The different metrics provided conflicting results. For the Shannon index, food restriction significantly reduced diversity (*p* = 0.030), while for observed OTUs, there was a significant change over time (*p* < 0.001), but no effect of Condition. The Inverse Simpson Index identified a Condition x Time interaction (*p* = 0.045). Plotting these metrics revealed visually similar results across them, wherein alpha diversity drops after injury and appears to recover over time for both groups, albeit with larger effect for food restricted rats. This suggests that there is truly an injury effect (“main effect of Time” given that all are injured), and potentially either a marginal interaction or main effect of Condition. While there is a larger drop in the Restricted rats, both groups recover by 14 days post-injury. These data are shown in Fig 6A.

The Bray-Curtis metric of beta diversity was analyzed with PERMANOVA (Condition*Time). There was a significant main effect of Condition (*p* = 0.007) and Time (*p* = 0.001), but the Condition x Time interaction did not reach significance (*p* = 0.051). These data are shown in Fig 6B.

Phyla proportions were analyzed with PERMANOVA (Condition*Time). There was no significant effect of Condition (*p* = 0.075) or Condition x Time (*p* = 0.498), however there was a main effect of Time (*p* = 0.001), suggesting an overall injury effect as phyla shifted then recovered. After multiple correction comparison, there was a significant shift from Day 1 to Day 3 for Firmicutes (decreased), Bacteroidota (increased), Proteobacteria (increased), and Actinobacteria (increased). From Day 3 to Day 7, all except Actinobacteria significantly changed proportions again (reversing initial changes). There were no significant changes from Day 7 to 14. These data are shown in Fig 6C.

### Behavior-Predictive Modeling using Microbiome Data

Multiple regression models were employed in an exploratory analysis to better understand the relationship between the gut microbiome and behavior. All analyses were conducted using multiple sessions of stable optimal P2 choice data. Prior to injury (4 sessions of behavior, pre-injury microbiome time point), LMER models including alpha diversity, Firmicutes, Bacteroidota, Firmicutes/Bacteroidota Ratio, Proteobacteria and their various combinations were evaluated in how well they accounted for optimal P2 choice. The best-fitting model included Firmicutes, Bacteroidota, and Proteobacteria. Though this was better than an intercept-only model, none were significant predictors, and the fixed portion of the model only explained 4% of the total variance in behavior. Because of the high subject-level variability, this analysis was repeated with no random effects which identified the intercept-only (mean value) model as the best fit.

To determine if alpha diversity or any of the individual phyla (or Firmicutes/Bacteroidota ratio) was predictive of chronic outcome on their own, we analyzed the final two weeks of behavior before the condition reversal (weeks 7-8 post-injury, 10 sessions). Each microbiome measurement time point was analyzed as the predictor with optimal P2 choice as the outcome. Only alpha diversity at 14D was predictive of long-term outcomes (*p* = 0.035), though pre-injury (*p* = 0.054) and 3D (*p* = 0.084) were near the threshold and accounted for between 11-16% of variance in behavior (Fig 7A). Interestingly, the one significant model (D14) indicated lower alpha diversity predicting better performance while the other near-significant models, the effect was in the opposite direction. This may be due to an interaction with injury or feeding condition. None of the individual bacteria levels were significant on their own.

**Figure 7.**
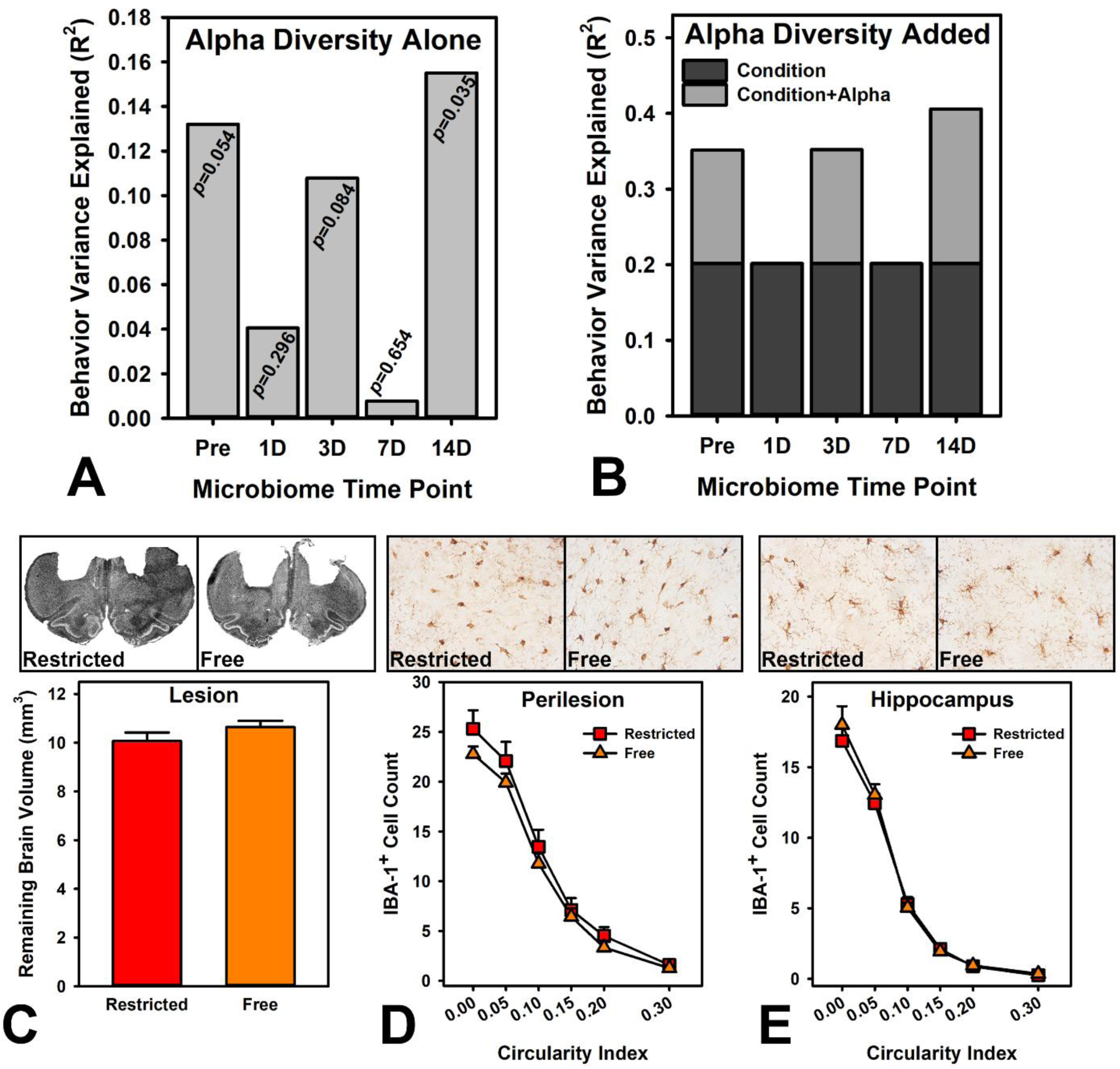
Model Comparisons and Histopathology. A) Alpha diversity (Shannon index) measured at the different time points was used as a predictor of chronic behavioral impairment. Only the 14-day post-injury time point was a significant predictor and accounted for 16% of variance. B) When adding alpha diversity (Shannon index) to Condition as a predictor, the pre-injury, Day 3, and Day 14 post-injury time points significantly improved the model, explaining an additional 15-21% of variance in the behavior. Days 1 and 7 were not improved by adding alpha diversity. C) There was no difference between the conditions in remaining brain volume. D) There was no difference between the groups in IBA-1 counts or morphology in the perilesion region. E) There was no difference between the groups in IBA-1 counts or morphology in the hippocampus.

Because our feeding manipulation affected the microbiome response to injury, we also performed model comparisons by assessing Condition, pre-injury P2 baseline, alpha diversity and all their additive combinations at each time point. Another set of models were compared which swapped individual bacterial phyla for diversity but were otherwise identical. Condition alone explained 20% of the variance and was the best model at 1D and 7D post-injury. However, alpha diversity improved model fit and variance explained at the pre-injury (+15% variance explained), 3D (+15% variance explained), and 14D (+21% variance explained) time points (Fig 7B). No individual bacteria levels improved model fits.

### Histology

There were no differences in total brain volume (*t*_(18.86)_ = 1.32, *p* = 0.204) between Conditions (Fig 7C). Total microglia counts were evaluated in the perilesion region and hippocampus using a Poisson LMER (fixed effects: Count ∼ Condition, random effects: Subject). There were no differences in either perilesion or hippocampus (*p* = 0.236, *p* = 0.496). We next evaluated whether the proportion of microglia that were more ameboid were different by Condition using a similar generalized LMER (Count ∼ Condition*Circularity). There were no differences in either perilesion or hippocampus (*p* = 0.244, *p* = 0.549; Fig 7D, E).

## Discussion

While psychiatric dysfunction is a common outcome for patients with TBI [5], the mechanisms driving disease and symptom development remain unclear. The gut microbiome causally modifies a variety of psychiatric diseases, including attention deficit hyperactivity disorder, bipolar disorder, and addictive disorders [10]. To effectively model relevant psychiatric-related behaviors in animals, food restriction is often required to motivate behavior which may have unanticipated effects on the gut microbiome and consequently, the target behavior. The current research indicated that this mild caloric restriction affected motivational and impulsivity measures on the RGT (Fig 3). Interestingly, caloric restriction only had minor effects on acquisition of optimal vs. suboptimal and risky decisions prior to the injury. TBI drove deficits for both groups in every measure (Figs 2, 3) except for impulsivity which was selective to the Restricted group. Moreover, caloric restriction amplified several of these deficits, including on the primary measure of interest, risk-based decision-making. Gut microbiome measurements of diversity and individual bacterial phyla indicated acute to subacute changes after TBI which were slightly amplified by caloric restriction (Fig 6). The diversity of the gut microbiome also predicted long-term impairments in decision-making (Fig 7A, B). These findings highlight a contributing role for the microbiome in the magnitude of post-injury deficits.

Because the current study required training before injury, some interesting comparisons can be made with literature focused solely on caloric restriction. Perhaps the most interesting finding from our pre-injury assessment was that there were no differences in decision-making or risk preference at baseline (Fig 2), despite minor changes in acquisition of those choice preferences. A prior study evaluated sensitivity to change in probabilities using a probability discounting task under multiple caloric restriction conditions [32]. They found that free feeding may reduce salience of consequences as indicated by an irregular discounting curve that included faster switching away from a degrading large option, but also higher maintenance of that choice through suboptimal probabilities. The lack of pre-injury decision-making differences in the current data stand in stark contrast to the other variables that were strongly affected with caloric restriction. Overall motivation to engage in the task was strongly reduced in the Free-fed condition, indicated by increased latencies and omitted trials as well as fewer trials completed (Fig 3), which is congruent with historical studies into the fundamentals of motivation. Likewise, free feeding also reduced the impulsive response (premature responding in the 5 s pre-choice phase; Fig 3A). Measuring a construct like impulsivity accurately in a rat may require some form of caloric restriction, what is commonly known as an “establishing operation” to motivate certain behaviors [18]. However, the primary decision-making variable of the RGT (optimal choice) may not require such an establishing operation, instead relying on the discriminative or comparative value between the different choices, at least for intact rats. These findings suggest some dissociation of *motivation to engage* versus *motivation to choose specific outcomes*.

While caloric restriction did not affect pre-injury decision-making, it significantly exacerbated the effect of injury (Fig 2). Further, caloric restriction also increased the effect of TBI on other variables (Fig 3), with the exception of reinforcer collection latency. While this may be somewhat concerning for broadly interpreting data requiring caloric restriction in rats after TBI, there were still robust injury effects in the Free-fed group in every single measure, except impulsive responding, as confirmed by comparison against baseline. Notably, the current findings of caloric restriction *worsening* outcomes after TBI go against multiple studies demonstrating protective effects of restriction. Acute deprivation (“fasting”) at time of injury spared cortex from a similar injury and reduced spatial learning impairments [33] and chronic restriction at 70% of free-feeding had similar benefits [34], even when instituted after injury [35]. This contradiction is interesting because prior studies used aversive conditions or negative reinforcement to motivate escape from a water maze, while the current study used a combination of positive reinforcement (sucrose pellets) and punishment (time out from earning) which are both directly related to the calorie restriction manipulation. This suggests that there may be some interactions of feeding and TBI that are specific to food-motivated behaviors. The reversal of feeding conditions allows for some inference surrounding caloric intake at time of injury vs. caloric restriction during testing. Flipping the feeding conditions had no effect on decision-making (Fig 4), suggesting that these effects may be more driven by conditions at the time of the injury or the long learning history and well-established preferences. In contrast, all non-decision variables significantly changed based on feeding condition (Fig 5), suggesting a similar effect of injury in both groups for these metrics that required caloric restriction to unmask. While reversing from restricted to free fed led to improvements in several metrics, this did not cause rats to “recover” anywhere close to the pre-injury baseline. Given the potential for caloric restriction to shape the gut microbiome, research requiring such restriction may necessitate further consideration of the role it plays in mediating behavioral outcomes.

Caloric restriction can drive strong gut microbiome alterations, particularly at extreme values (<75%) or due to intermittent feeding [19, 20]. The caloric restriction used in the current study placed rats at approximately 80-85% of free-feeding weight (Fig 1C). This is a relatively mild restriction and may more closely approximate real life compared to a free-fed laboratory rat. Thus, we might not anticipate drastic differences in gut microbiomes. Indeed, at the pre-injury time point, there were no gross differences in alpha diversity or any of the major phyla (Fig 6A, C). However, beta diversity, or the dissimilarity between subjects, slightly increased due to feeding condition (Fig 6B). Studies suggest that caloric restriction could function more as a catalyst for the microbiome, allowing rapid shifts in response to a challenge like illness [20, 21]. This appears consistent with the current results, where the dysbiosis was greatest or perhaps had the slowest recovery in the Restricted group, reflected by alpha diversity metrics. However, there were also common elements to the effect of TBI as seen in the similar shifts in beta diversity and bacterial phyla for both groups (Fig 6B-C). Notably, the gut microbiome largely recovered by 14 days after injury. This acute to subacute assessment was chosen based on prior data that the gut microbiome had recovered in rats after TBI by 30 days post injury [8], but that study was limited in its assessment with only a 3-day post-injury acute time point. One conclusion might be that the gut is not involved in chronic deficits because of this resolution compared to behavior. However, there may be a role for the gut microbiome in the acute and subacute period that sets up long-term functional impairment during this critical time of brain recovery and remodeling.

In a multitude of psychiatric disorders, the gut microbiome has a bidirectional relationship with symptoms and pathology. As an example, chronic unpredictable stress reduced microbiome diversity in rats while inducing anhedonia representing an environment to symptom and gut effect [36]. However, a fecal transfer from patients with clinical depression into microbiome-depleted rats was sufficient to induce anxiety and depressive symptoms [37] representing a gut to symptom process. Together these highlight the truly bidirectional nature of the gut microbiome. It is possible that microbiome dysbiosis from TBI may feed forward and augment injury symptoms. Though caution should be taken not to over-interpret correlative data, this relationship can be explored using the current dataset. By comparing the degree to which gut dysbiosis predicts long-term outcomes in optimal decision-making, we can set up targeted hypotheses for future experiments. When evaluating alpha diversity on its own, without any other predictors, the day 14 measurement was the only significant predictor of long-term impairment. However, the current data may be somewhat limited in ranges of diversity and optimal decisions (both impacted by TBI) given the lack of a sham control condition. Indeed, when we considered other factors such as pre-injury baseline (change from baseline = “injury effect”) or the feeding condition which clearly affects outcomes, alpha diversity became more important in accounting for variance in outcome. Alpha diversity at 3- and 14-days post-injury improved predictive power (adding 15-21% of variance explained; Fig 7B). Moreover, even the pre-injury time point alpha diversity measurement added explanatory power (+15% variance explained), suggesting an early interaction. Together, these suggest that diversity is not the one driver on its own, but likely interacts with other causal variables. While these should be interpreted with caution as they are exploratory analyses, they indicate a causal role for the gut microbiome in modifying long-term outcomes from TBI which is supported by other studies using more direct manipulations [6, 15].

The mechanism by which the gut microbiome contributes to long-term deficits from TBI is not resolved by the current data. Two major, and potentially complementary, hypotheses are that peripheral inflammation augments the neuroinflammatory process from TBI or that neurotransmitter disruption stems from vagal dysregulation. Our data indicate that the phylum of proteobacteria, which contains many pro-inflammatory bacteria, drastically increases within 3 days post-injury (Fig 6C). However, no models that included proteobacteria were significant predictors of function, weakening some of this support. Moreover, we did not observe gross changes in microglia number or morphology between groups (Fig 7D, F). These data do not exclude this hypothesis but indicate more refined measurements may be needed to further evaluate. Vagal dysregulation of neurotransmitter systems, particularly serotonin, are indicated in other models of psychiatric disease and the gut microbiome [10, 38]. This has not been directly evaluated in TBI, but older studies in animals indicated vagal nerve stimulation may be efficacious in treating TBI [39] and recent human data preceding clinical trials show promise for the ability to apply this transcutaneously [40, 41]. More data will be needed to elucidate the specifics by which vagal stimulation, the gut microbiome, and neurotransmitter function interact to promote recovery from TBI.

In summary, the current study establishes novel findings for caloric restriction and decision-making with a potential role for the gut microbiome as a mediator. Interestingly, in the absence of injury, restriction did not substantially change probability-based decisions. However, caloric restriction at the time of injury worsened injury effects, which contrasts with several prior TBI studies that did not use food-reinforced behavioral assessment [33–35]. Moreover, we confirmed that several behavioral variables were partially motivation-dependent on restriction, and that important variables to TBI like impulse control may require caloric restriction to measure. Finally, TBI induced gut dysbiosis for both groups, and this was predictive of long-term behavioral outcomes, particularly when accounting for other factors. These data indicate a need to further dissociate this mechanism to determine if it may be a useful route for effective TBI therapies.

### Transparency, Rigor, and Reproducibility

One exclusion was made based on little damage to the cortex, suggesting a surgical mistake in the injury. Full code for all analyses (R script) will be made available. Detailed tables in the supplement provide all statistical values, including subject-level random effect information.

Individuals were nominally blinded to rat condition, but with caloric restriction vs. free feeding, it may be obvious during behavior testing which rats were in which condition. However, all testing is performed with automated computer-controlled operant chambers, mitigating concerns over bias in this. All other analyses were performed blind.

## Data Availability Statement

Data will be made publicly available at the Open Data Commons for Traumatic Brain Injury (search Vonder Haar). Associated analysis code will be publicly posted to the corresponding author’s Github (https://github.com/VonderHaarLab/).

## Author Contributions

Reagan Speas: Designed research, performed research, analyzed data, wrote manuscript. Jenna McCloskey: Designed research, performed research, analyzed data, wrote manuscript. Noah Bressler: Performed research. Michelle Frankot: Analyzed data. Carissa Gratzol: Performed research. Kris Martens: Designed research, performed research, wrote manuscript. Cole Vonder Haar: Designed research, analyzed data, wrote manuscript.

## Acknowledgements

We would like to thank Sarah Wampler and the other members of the Injury and Recovery Laboratory for helping with behavioral testing. This research was funded in part by seed awards from the Chronic Brain Injury Program and Neuroscience Research Institute at Ohio State University.

## Supplemental Results

The following decision-making variables were analyzed, but not included in the main manuscript: Overall score on the RGT (beneficial vs. detrimental choices: [P1+P2] / [P3+P4]), likelihood of staying with a choice (p(Stay)), likelihood of staying with a choice after a win (p(Stay|Win)), likelihood of staying with a choice after a loss (p(Stay|Loss)). Full statistics can be found in the tables below alongside other variables.

There were differences in acquisition between the groups on the Score, p(Stay), and p(Stay|Win), but not p(Stay|Loss). However, when compared at baseline, there were no differences between the groups on any measures (Fig S1, left of line break).

After injury, there were group differences in Score, p(Stay) and p(Stay|Win), but not p(Stay|Loss). However, a within-subjects comparison confirmed an injury effect for each of the variables (Fig S1, right of line break).

After the condition reversal, there was a significant overall decrease in Win-Stay likelihood for the (formerly) Restricted group (*p* = 0.044), but no difference on other measures (Fig S2).

All statistical analysis summaries are in Supplement 2, an .xlsx file. Each tab contains sets of analyses from acquisition, post-injury, post-reversal, and microbiome data. Full datasets will be made available on ODC-TBI and all analysis code made available on the author’s Github (https://github.com/VonderHaarLab/).

**Fig S1.**
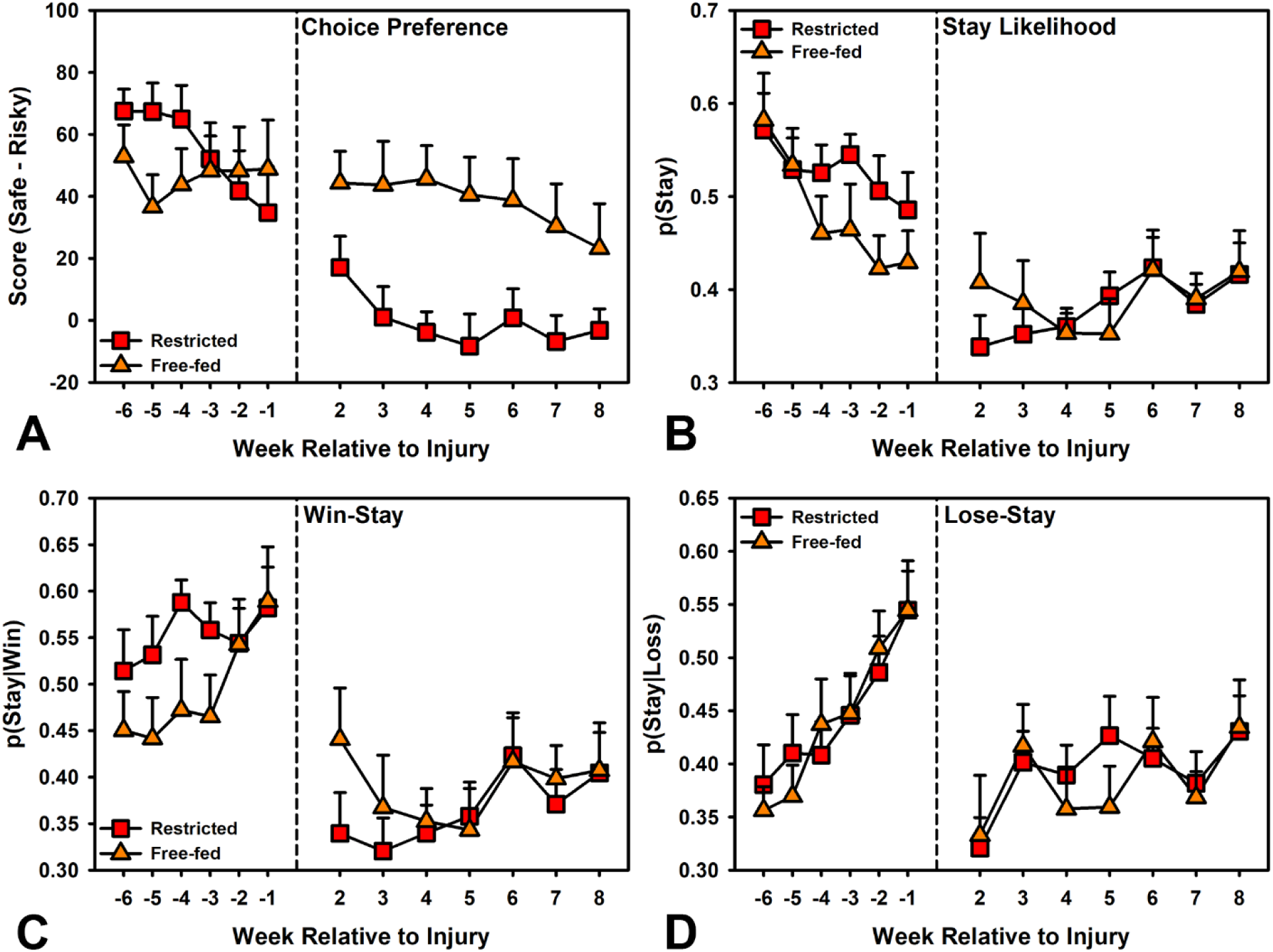
Supplemental RGT decision-making variables during acquisition and post-injury. There were significant group differences in rate of acquisition for panels A, B, and C (left of line break). Comparison of just the baseline (-1 weeks relative to injury) revealed no differences. After injury, there were group differences in panels A, B, and C (right of line break).

**Fig S2.**
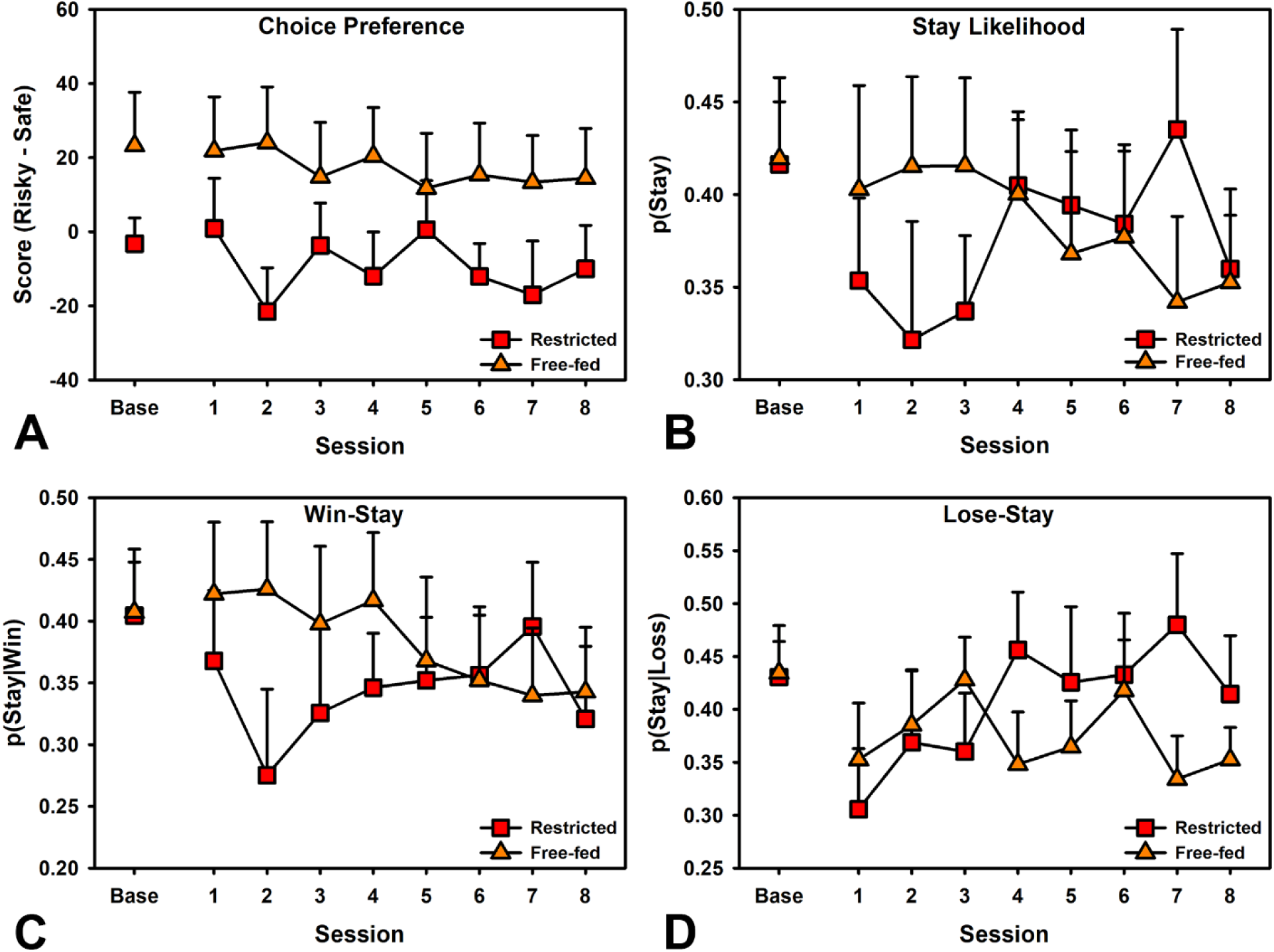
Supplemental RGT decision-making variables after condition reversal. The condition reversal only significantly affected panel C (*p* = 0.044).

